# PRCnet: An Efficient Model for Automatic Detection of Brain Tumor in MRI Images

**DOI:** 10.1101/2023.09.28.560042

**Authors:** Ahmeed Suliman Farhan, Muhammad Khalid, Umar Manzoor

## Abstract

Brain tumors are the most prevalent and life-threatening cancer; an early and accurate diagnosis of brain tumors increases the chances of patient survival and treatment planning. However, manual tumor detection is a complex, cumbersome and time-consuming task and is prone to errors, which relies on the radiologist’s experience. As a result, the development of accurate and automatic system for tumor detection is critical. In this paper, we proposed a new model called Parallel Residual Convolutional Network (PRCnet) model to classify brain tumors from Magnetic Resonance Imaging. The PCRnet model uses several techniques (such as filters of different sizes with parallel layers, connections between layers, batch normalization layer, and ReLU) and dropout layer to overcome the over-fitting problem, for achieving accurate and automatic classification of brain tumors. The PRCnet model is trained and tested on two different datasets and obtained an accuracy of 94.77% and 97.1% for dataset A and dataset B, respectively which is way better as compared to the state-of-the-art models.

## Introduction

The human brain is the most complicated organ of the human body [1]. It is responsible for controlling the entire nervous system [2]. Due to its intricate design, even a slight malfunction in the brain cells can cause severe disruption in the performance of connected organs [3]. A brain tumor is a disease that occurs due to the abnormal growth of brain cells [4]. This uncontrolled growth of cells poses severe problems to human health [5]. It is one of the leading causes of death in adults and children today [6]. Brain tumors also come in different forms, and their symptoms are likely to vary depending on the type of tumor [7]. Brain tumors are the world’s most common type of cancer [8]. Even though brain tumors comprise only 2% of all cancers, they are disproportionately to blame for cancer-related fatalities [9].Brain tumors can broadly be classified into two main categories: primary tumor and secondary tumor [10]. Primary tumors, which account for 70% of all brain tumors, develop from the brain’s own cells. Secondary tumors, comprising the remaining 30%, are tumors that begin somewhere else in the body and then spread to the brain through the bloodstream [11]. In most cases, primary brain tumors are benign, while secondary brain tumors are usually more malignant [12].

The tumor location and growth rate can affect the nervous system functionality differently and may cause headaches, seizures, memory loss, balance problems, etc [3]. There are two types of brain tumors namely malignant and benign tumors [13]. Malignant tumors are considered to be the most perilous. They tend to grow rapidly, infiltrate surrounding tissue aggressively, spread to other areas in the brain, and can cause severe damage to the nervous system [14]. On the other hand, benign tumors tend to grow at a slower pace [15].In order to identify the disease, an individual must undergo a series of neurological examinations which assess their vision, balance, hearing, strength, and other functions. If any anomalies are detected during these assessments, the next step is to conduct either a brain Computerised Tomography (CT) or Magnetic Resonance Imaging (MRI) scan [3]. MRI is widely used as a non-invasive diagnostic tool for detecting various types of brain tumors and provides detailed information on the brain tissues [16]. Also, It is one of the best and most accurate tests for tumor identification due to its ability to produce high-quality images of human anatomy in multiple dimensions [17]. MRI uses radio waves and magnetic resonance to retrieve detailed brain information and can help doctors in tumor diagnosis [18]. The brain can be imaged in three planes, coronal, axial and sagittal - see Figure 1 and multi-angle images can be obtained [19].

**Fig 1.**
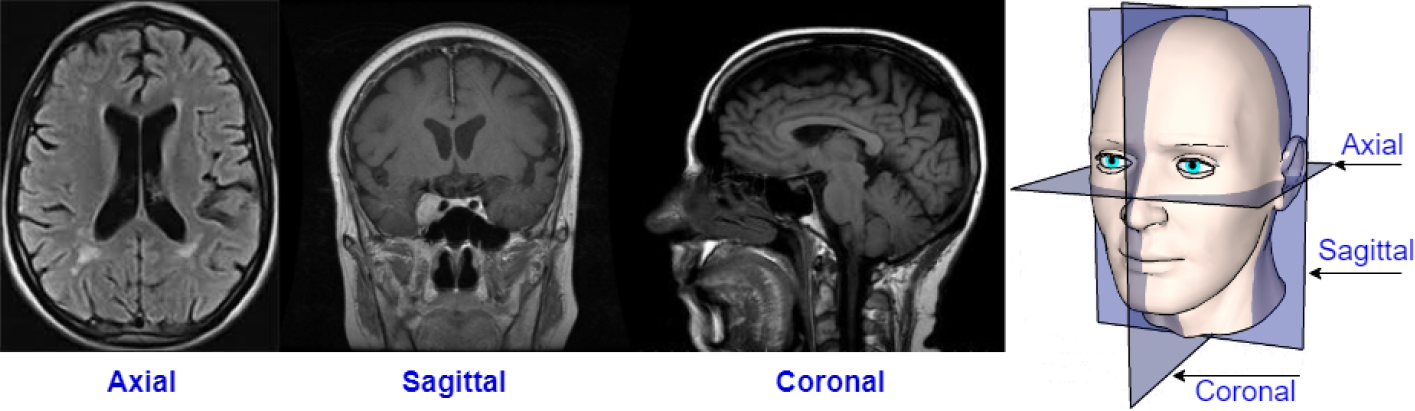
MRI Image Planes.

Early detection of tumors is crucial in identifying the severity of the brain malformation, which is essential for deciding the most appropriate treatment plan [20]. In addition, accurately identifying the severity of a brain malformation can assist medical professionals in making informed decisions related to treatment options [21]. Also, it increases chance of patient survival is and improved quality of treatment [22]. The task of manually detecting tumors is arduous, intricate, and time-consuming, and may often be erroneous owing to the diverse nature of cancer cells. Additionally, it relies on the radiologist’s expertise [23]. As a result, it is imperative to develop and dependable automated tumor detection and segmentation technique that can effectively assist radiologists with prompt and efficient classification [24].

Medical image processing is critical in assisting healthcare professionals in diagnosing a wide range of medical conditions [25]. In conjunction with magnetic resonance imaging (MRI), biomedical image processing facilitates the detection and segmentation of brain tumors [26].Deep neural networks have emerged as a powerful tool for processing medical images, achieving remarkable results in various applications [27]. In addition, the use of deep neural networks for detecting brain tumors from MRI scan images has become increasingly popular due to their high accuracy [28]. Convolutional Neural Network (CNN) is one of the most effective deep learning designs for learning representations from an input signal with different levels of abstraction [29]. CNN is an attractive technology with many advantages over traditional machine learning algorithms. The benefits include a) increased accuracy and efficacy over raw spectra (thus there is no need to process the data prior to classification) [30], and b) infer hidden patterns from complex data and images [31]. Using an intelligent, fast and error-free system for detecting brain tumors using MRI images is very important to help clinicians accurately detect the tumors [32].

The CNN has performed exceptionally well in image classification tasks and has achieved better performance than rival human experts in classifying medical images [29]. For example, Aurna et al. proposed a new approach by using a two-stage feature ensemble of CNN to classify the brain tumors. By experimenting and testing the models (VGG19, EfficientNetB0, InceptionV3, ResNet50, Xception, and Scratched CNN) on three datasets of MRI images, the models EfficientNetB0, ResNet50, and Scratched CNN are selected for the first stage of the ensemble. The two models with the highest accuracy are selected for the second stage of the ensemble. The proposed model are trained and tested on three databases and the fourth dataset resulted from merging the three datasets, and achieved average accuracy of 99.13% [33]. Although the accuracy obtained is very good, however, the proposed model when tested on dataset A and B, obtained 92.59% and 94.2% accuracy respectively. This difference in results is due to difference in 1) datasets and 2) datasets splitting into groups for testing, validation, and training. Furthermore, the model is considered very complex because it combines three models, two of which are designed to classify a thousand classes from the Imagenet dataset [34].

In this paper, we proposed the state of the art model called PRCnet, to overcome the limitations and problems that exist in the literature. The existing models are either very simple (obtain good accuracy on very small dataset, but under-perform on large dataset) or very complex (incorporate a number of models that are essentially not designed to handle medical images). All this motivated us to develop a new model that uses parallel layers with different filter sizes and connections between layers to provide excellent accuracy in classifying brain tumors. The main contributions of the PRCnet model are:

- A novel model called PRCnet for the detection of brain tumors in MRI images.
- Computationally less expensive where total parameters for PRCnet model are 21413060 as compared to 144853060, 35533911, 134276932 for Musallam et al. [35], Aurna et al. [33], and VGG16 [36] respectively.
- Different filter sizes used with parallel layers help the model to recognise small and large features which leads to improved accuracy.
- A comprehensive comparison is performed with state-of-the-art models, the PRCnet model achieved an accuracy of 97.1 and 94.77 for datasets A and B respectively.
- The PRCnet robustness is validated by using a cross-validation technique on datasets A and B.

The Figure 2 shows a summary of the proposed methodology. In the first stage, the dataset of brain MRI images used for training and testing is determined namely dataset A and B. The second stage is data pre-processing (such as image resizing), afterwards, the images are divided into three training, validation, and testing sets. The third stage is to design the PRCnet model, and train / test it on datasets A and B. The fourth stage is to identify some of the most popular and pre-trained standard models (such as VGG16, VGG19 and MobileNetV2) and use the learning transfer from the ImageNet dataset, then train/test the models on dataset A and B. Moreover, we train and test some state-of-the-art relevant papers on dataset A and B. Finally, we compared the results of the PRCnet model with the results of the (VGG16, VGG19, MobileNetV2, Aurna [33], Chattopadhyay [37] and Musallam [35]) models.

**Fig 2.**
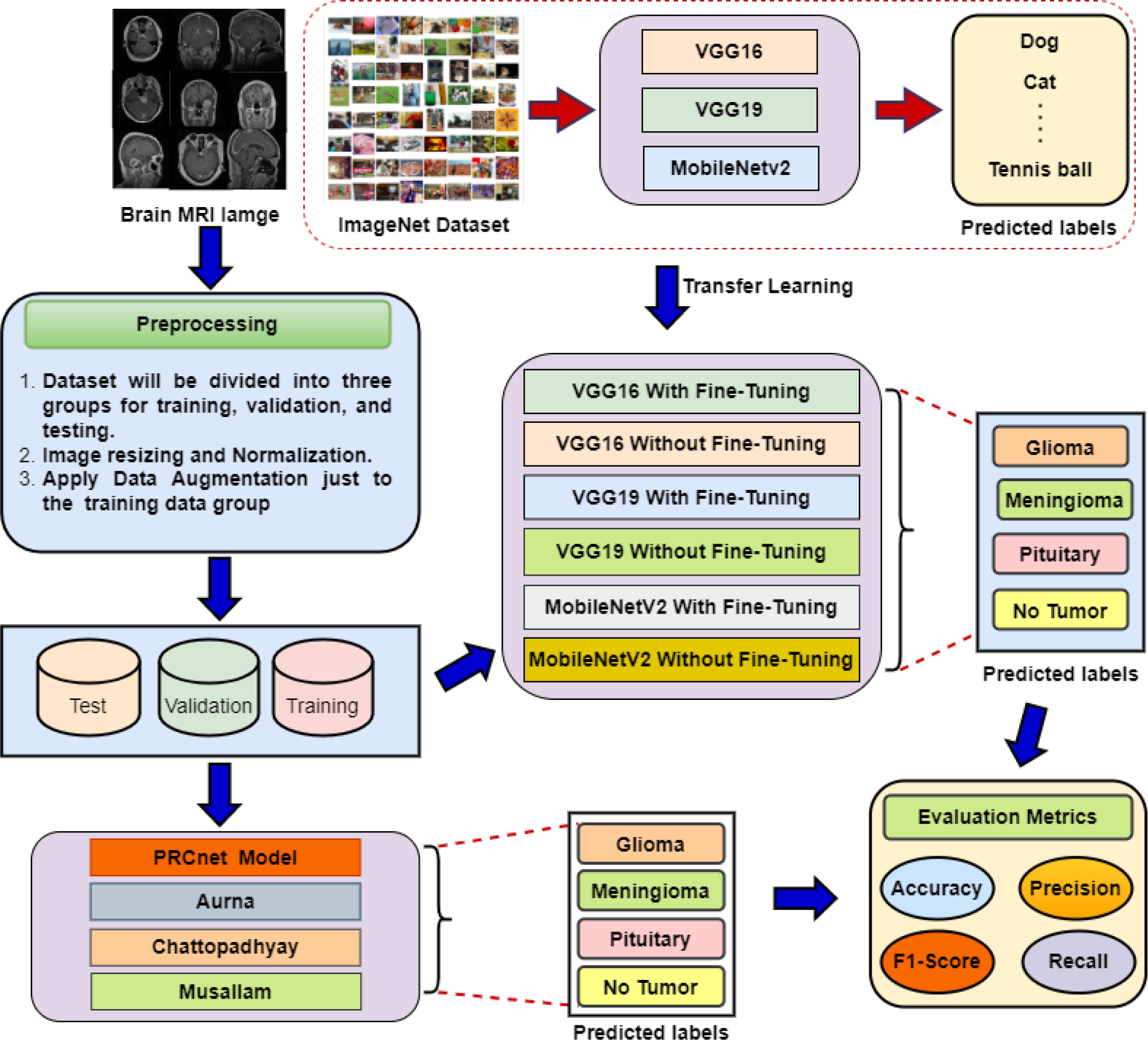
Flow of various steps including pre-processing, transfer learning, feature extraction, classification, and evaluation matrix.

## Literature review

Researchers have presented many studies to diagnose brain tumors using deep learning techniques in recent years by using MRI Image as input to the deep learning model.

Applying these methods to medical signals and images can aid non-specialist physicians in making proper clinical decisions.

Aurna et al. [33] proposed a new approach by using two-stage feature ensemble of CNN to classify the brain tumors. By experiment and testing the models (VGG19, EfficientNetB0, InceptionV3, ResNet50, Xception, and Scratched CNN) on three datasets of MRI images, the models EfficientNetB0, ResNet50, and Scratched CNN are selected for the first stage of the ensemble. The two models with the highest accuracy are selected for the second stage of the ensemble. The proposed model is trained and tested on three databases and the fourth dataset resulted from merging the three datasets, and achieved average accuracy of 99.13%.

Rai and Chatterjee [38] proposed the CNN model (LeUNet), which is less complex, has fewer layers, and faster processing time for detecting brain tumors. The proposed model is simulated on a dataset containing 253 images and compared with VGG-16, U-Net, and Le-Net, where the accuracy of proposed model is 98% and 94% on cropped and uncropped images respectively, and a speedy processing time of only 252.36 seconds. However, the model is not tested on a larger database as well as it did not classify the types of brain tumors

Masood et al. [39] proposed brain tumor segmentation and precise classification using custom Mask Region-based CNN (Mask-RCNN) with a dense-net-41 backbone architecture trained by transfer learning. Two benchmark datasets are used to evaluate Mask-RCNN and the results show that the Mask-RCNN can accurately detect tumor and precise tumor locations. The model achieved an accuracy of 98.34% for classification and 96.3% for segmentation, however, the model is not tested on a large dataset to analyse the model robustness. The model is tested on two datasets, the first dataset contained 3064 MRI images whereas the second one contained only 253 MRI images, which are not sufficient to demonstrate the robustness of the model.

Abd El Kader et al. [40] proposed a differential deep convolutional neural network model that uses magnetic resonance images to classify different types of brain tumors. The authors used CNN architecture’s differential operators to create extra differential feature maps in the original CNN feature maps to improve the proposed model. This enables model to analyze the pixel orientation pattern based on contrast calculations and classify large number of images with high accuracy. A dataset of 25,000 MRI images collected from Tianjin Universal Center of Medical Imaging and Diagnostic (TUCMD) is used for training and testing. The model achieved 99.25% accuracy, 95.23% F-score, 95.89% sensitivity, 97.22% precision, and 93.75% specificity. However, the model is not evaluated on a dataset that contains more than two classes (tumor types).

Hapsari et al. [41] proposed deep learning model based on VGG-16 namely en-CNN to overcome the hyperparameter tuning problem by using hyperparameters optimizer on top of existing VGG-16 layers. The proposed approach does not require a prior segmentation process for the purpose of classification but rather classifies brain tumors directly. The BraTS 2018 dataset [42] containing four MRI sequences, FLAIR, T1, T2 and T1CE, is used for training and testing, the results shows that the proposed method achieved an accuracy of 95.5%, 95.5%, 94% and 97% for T1, T1CE, T2 and FLAIR respectively. However, the model doesn’t classify the tumor types and also classifies T1, T1CE, T2, and FLAIR separately.

Kokila et al. [43] proposed a model based on the convolutional neural network to detect and classify tumors, instead of using the individual model for each classification task, a multi-task classification model is used to classify the MRI images. This method divides and locates the brain tumor and is suitable for classifying thrush type tumors. The model is tested on dataset collected from different sources and achieved 92% accuracy. However, model robustness analysis and detail results for each dataset is missing.

Irmak [44] proposed three CNN models for three different classification tasks, the first one classify tumor presence, the second one classify tumor into five types (metastatic, normal, meningioma, glioma and pituitary), and finally, the third one classify the tumor in three grades (Grade II, Grade III and Grade IV). The proposed models achieved 99.33%, 92.66% and 98.14% accuracy respectively. The accuracy is high for tumor detection, however, accuracy decrease significantly for tumor classification.

Gu et al [45] proposed a model using convolutional dictionary learning with local constraints (CDLLC) for classifying brain tumors. Multi-layered dictionary learning is combined with convolutional neural network architecture, and two multi-category classification tasks were designed based on the Cheng and REMBRANDT datasets to test and train the proposed method. The results showed that the method effectively classifies brain tumors based on magnetic resonance images, however, the dataset used for training and testing is relatively very small (only 1000 images) which is insufficient to test model robustness.

DíIaz-Pernas et al. [46] proposed a model based on multiscale convolutional neural network for automatic tumor segmentation and classification. The input images are processed at three spatial scales along with different processing paths. The proposed model is trained and tested on a database containing 3064 MRI images, including three types of tumors (glioma, meningioma and pituitary tumor). The method achieved an accuracy of 97.3% in classifying brain tumors, also obtained distinct segmentation performance, an average of 0.940, 0.828 and 0.967 for sensitivity, Dice index and pttas value respectively. However, the model lacks durability analysis and testing on a larger dataset.

Noreen et al. [47] suggested a model of multi-level features extraction for brain tumors identification. This study presented two various scenarios. In the first scenario, features from various DensNet blocks are extracted from a trained DensNet201 model and passed to classify brain tumors using a softmax classifier. In the second scenario, the pre-trained Inceptionv3 model is used to extract features from the different Inception modules, which are passed to a softmax to diagnose the brain tumor. Publicly available tumor dataset is used to evaluate both scenarios, the results of the accuracy-test is 99.34% for Inception-v3 and 99.51% for DensNet20. The model has not been tested on a large database to evaluate the model robustness.

Rajinikanth et al. [48] developed a deep learning architecture (DLA) for the purpose of automated detection of brain tumors using magnetic resonance imaging. To discover brain diseases, they (i) suggested implementing a pre-trained deep learning structure such as (ResNet101, AlexNet, ResNet50, VGG19 and VGG16) and classification using SoftMax. (ii) using a pre-trained deep learning architecture but a decision tree, SVM-linear, SVM-RBF and k nearest neighbour (KNN) based on in-depth features for classification; (iii) and custom VGG19 with sequentially embedded features and other handcrafted features to improve accuracy. The results proved that VGG19 achieved better results compared with (VGG16, ResNet101, AlexNet, and ResNet50), where the VGG19 accuracy is of 97%, 98% 99% for the modalities T1C, T2, and Flair respectively. The study is limited to tumor detection and doesn’t deal with the classification of tumor type.

Musallam et al. [35] proposed three preprocessing steps to improve the MRI brain images, such as removing the confusing variables, denoising the MRI images, and enhancing the contrast of these images. In addition, a new Deep Convolutional Neural Network (DCNN) architecture is proposed to effectively diagnose three types of brain tumor (glioma, meningioma, and pituitary). The model is trained and tested on a dataset consisting of 3,394 images, and the average accuracy is 98.22%. Finally, the proposed model results are compared with well-known techniques such as VGG16, VGG19, and CNN-SVM.

Fayaz et al. [49] proposed a model to do the binary classification of brain MRI. Firstly, the grayscale MRI images are converted into RGB images. Secondly, discrete wavelet transform (DWT) is used to make feature extraction. Thirdly, statistical technique for each color channel such as (energy, variance, kurtosis, correlation, entropy, contrast, mean, skewness, and homogeneity) is used for feature reduction. Finally, blended artificial neural networks are used to perform the classification by applying the concept of majority voting between results from each ANN for each color channel.

Rizwan et al. [50] proposed Gaussian Convolutional Neural Network (GCNN) for brain tumor classification and used many pre-processing filters to enhance the classification task. The proposed method is trained and tested on two datasets. The first dataset is brain tumor dataset [51] and the second dataset from “The Cancer Imaging Archive (TCIA)” [52]. The proposed method achieved an accuracy of 99.8% and 97.14% on both datasets respectively.

Chattopadhyay and Maitra [37] proposed a CNN model for detecting brain tumors from magnetic resonance imaging. The model consists of two Convolution layers with activation function Relu. Each Convolution layer follows a batch normalization layer and a maxpooling layer. The model has two fully connected layers, the first layer has the ReLU activation function whereas the second one has a Softmax activation function. The model is trained and tested on the BraTS 2020 dataset, and achieved 99.74% accuracy.

Raza et al. [53] proposed DeepTumorNet for brain tumor classification. The DeepTumorNet is a hybrid deep learning model based on the GoogLeNet model after removing the last five layers and adding 15 new layers instead of eliminating ones. they used a brain tumor dataset [51] to train and test the proposed model. The results were 99.67%, 99.6%, 100%, and 99.66% for accuracy, precision, recall, and F1-score, respectively.

Jun and Liyuan [54] developed a new model for classifying brain tumors using magnetic resonance imaging. The proposed model integrates an attention mechanism focusing on important information with a multipath network to improve accuracy. The model achieved an accuracy of 98.61% when it was tested using a brain tumor dataset [51].

Several deep learning have been proposed for tumor detection and classification with satisfactory results as shown in the table 1, however, most of the proposed models are trained and tested on a small dataset which is insufficient to evaluate model unbiased robustness and accuracy. To compensate for the lack of data, some researchers develop their models based on standard models such as (VGG16) as these models are pre-trained on a huge database and are designed to classify thousands of images. However, these strategies are associated with more computational costs. Moreover, some models are limited to tumor detection and does not classify tumor types. In general, the existing models lacks features that make the model more generalised and give acceptable results through testing on two or more datasets with cross-validation. Therefore, we designed a generalised PRCnet model by performing many experiments to choose the optimal filter size and kernel size for each convolutional layer. We analyzed that using different filter kernel sizes with parallel layers increased the model’s accuracy. Also, we used connections between layers to obtain features from different levels and address the problem of gradient vanishing. Furthermore, a dropout layer is used to help overcome the problem of overfitting.

**Table 1.** Summary review of recent developments in brain tumor classification.

| Reference | Summary | Dataset | Result |
| --- | --- | --- | --- |
| Aurna et al. [33] | Proposed a new approach by using two-stage feature ensemble of CNN to classify the brain tumors. | Dataset 1 contains 3064 MRI brain images [51]. Dataset 2 contains 3264 MRI brain images [55]. Dataset 3 contains 4292 MRI brain images [56]. | The average accuracy was 99.13% |
| Rai and Chatterjee [38] | Proposed the CNN model (LeUNet), which is less complex, has fewer layers, and faster processing time for detecting brain tumors. | Brain MRI Images for Brain Tumor Detection [57]. | The accuracy was 98% and 94% on cropped and uncropped images, respectively. |
| Masood et al. [39] | proposed brain tumor segmentation and precise classification using custom Mask Region-based CNN (Mask-RCNN) with a dense-net-41 backbone architecture trained by transfer learning. | Brain tumor dataset [51]. Brain MRI Images for Brain Tumor Detection [57]. | The accuracy is 98.34% and 96.3% for classification and segmentation, respectively. |
| Abd El Kader et al. [40] | proposed a differential deep convolutional neural network model that uses magnetic resonance images to classify different types of brain tumors. | A dataset of 25,000 MRI images collected from Tianjin Universal Center of Medical Imaging and Diagnostic (TUCMD). | The model achieved 99.25% accuracy, 95.23% F-score, 95.89% sensitivity, 97.22% precision, and 93.75% specificity. |
| Hapsari et al. [41] | proposed deep learning model based on VGG-16 namely en-CNN to overcome the hyperparameter tuning problem by using hyperparameters optimizer on top of existing VGG-16 layers. | The BraTS 2018 dataset [42]. | accuracy of 95.5%, 95.5%, 94% and 97% for T1, T1CE, T2 and FLAIR, respectively. |
| Kokila et al. [43] | proposed a model based on the convolutional neural network to detect and classify tumors, instead of using individual model for each classification task, a multi-task classification model is used to classify the MRI images. | The model was tested on dataset collected from different sources. | Achieved 92% accuracy. |
| Irmak [44] | proposed three CNN models for three different classification tasks, the first one classify tumor presence, the second one classify tumor into five types (metastatic, normal, meningioma, glioma and pituitary), and finally, the third one classify the tumor in three grades (Grade II, Grade III and Grade IV). | 1- The reference image database to evaluate therapy response (RIDER) [58].<br>2- The Repository of Molecular Brain Neoplasia Data (REMBRANDT) [59].<br>3- The cancer genome atlas low-grade glioma (TCGA-LGG) [60].<br>4- Brain tumor dataset [51] | Achieved 99.33%, 92.66% and 98.14% accuracy, respectively. |
| Gu et al. [45] | proposed a model using convolutional dictionary learning with local constraints (CDLLC) for classifying brain tumors. | 1- REMBRANDT [59]. 2- Brain tumor dataset [51] | The average accuracy is 97.64% on REMBRANDT dataset, 996.34% on Brain tumor dataset. |
| Díaz-Pernas et al. [46] | proposed a model based on multiscale convolutional neural network for automatic tumor segmentation and classification. The input images are processed at three spatial scales along different processing paths. | Brain tumor dataset [51] | The method achieved an accuracy of 0.973 in classifying brain tumors.<br>The method also obtained a distinct segmentation performance, an average of 0.940, 0.828 and 0.967 for sensitivity, Dice index and pttas value, respectively. |
| Noreen et al. [47] | Suggested a Model of multi-level features extraction for brain tumors identification. | Brain tumor dataset [51] | The accuracy were 99.34% for Inception-v3 and 99.51% for DensNet20 . |
| Rajinikanth et al. [48] | Developed a deep learning architecture (DLA) for the purpose of automated detection of brain tumors using magnetic resonance imaging. | BRATS [42], and The Cancer Imaging Archive (TCIA) | The results proved that VGG19 achieved better results compared with (VGG16, ResNet101, AlexNet, and ResNet50), where the VGG19 accuracy is of 97%, 98% 99% for the modalities T1C, T2, and Flair, respectively. |
| Musallam et al. [35] | proposed three preprocessing steps to improve the MRI brain images, such as removing the confusing variables, denoising the MRI images, and enhancing the contrast of these images. In addition, they proposed a new Deep Convolutional Neural Network (DCNN) architecture. | 1- Brain Tumor Classification (MRI) [55]. 2-Brain MRI Images for Brain Tumor Detection [57]. | The average accuracy was 98.22%. |
| Fayaz et al. [49] | pproposed a model to do the binary classification of brain MRI. |  | Achieved 98% accuracy. |
| Rizwan et al. [50] | proposed Gaussian Convolutional Neural Network (GCNN) for brain tumor classification and used many pre-process filters to enhance the classification task. | Brain tumor dataset [51] . 2- The Cancer Imaging Archive (TCIA) [52]. | The accuracy of 99.8% and 97.14% on both datasets, respectively . |
| Chattopadhyay and Maitra [37] | Proposed a CNN model for detecting brain tumors from magnetic resonance imaging. | BraTS 2020 dataset [42] . | The accuracy achieved was 99.74. |
| Raza et al. [53] | proposed DeepTumorNet for brain tumor classification. The DeepTumorNet is a hybrid deep learning model based on the GoogLeNet model after removing the last five layers and adding 15 new layers instead of eliminating ones. | Brain tumor dataset [51] | The results were 99.67%, 99.6%, 100%, and 99.66% for accuracy, precision, recall, and F1-score, respectively. |
| Jun and Liyuan [54] | developed a new model for classifying brain tumors using magnetic resonance imaging. The proposed model integrates an attention mechanism focusing on important information with a multipath network to improve accuracy. | Brain tumor dataset [51] | The model achieved an accuracy of 98.61%. |

## Convolution Neural network Overview

Neural networks are powerful techniques capable of handling large amounts of data with high accuracy. The deep neural architectures has brought advancement in ANN such as Recurrent Neural Network (RNN) and CNN. Deep neural networks help understand and analyse hidden patterns from data and images [31]. It is excellent in extracting features from images [61]. Therefore, it is used for detection, image segmentation, registration, localisation and classification [62]. The CNN is very commonly used in medical image analysis [63]. The deep learning approach has recently achieved promising results in analysing medical images [64]. A CNN can automatically represent features from training data and update parameters for convolutional filters during training [65]. The convolutional neural network architecture consists of several layers, such as an input layer, hidden layers, and an output layer [66]. A hidden layer may contain several different layers, such as a pooling layer, Fully-connected layers, a Convolutional layer, or an Activation Function. Figure 3 shows a general CNN architecture for a brain tumor classification task. In MRI images, the image is fed into the network, several stages of convolution layers, activation functions, and pooling layers follow. After that, a layer or more of the Fully-connected layers and then the classification layer (Output layer) is processed. This is the most common architecture for classifying image models. However, in recent years, many models and architectural changes have been proposed to improve classification accuracy and reduce computation costs such as VGG [36], ResNet [67] and MobileNetv2 [68]. Next, we will present out model for improving the accuracy of tumor detection in MRI images and then it will compared with the state of the art solutions.

**Fig 3.**
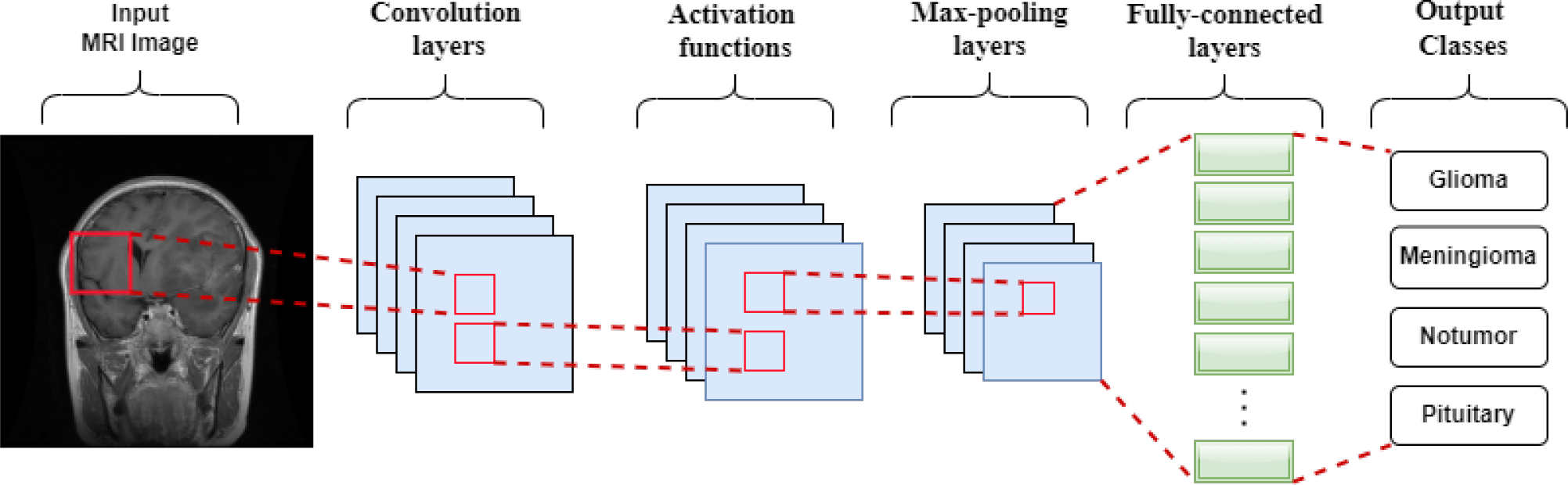
A CNN architecture for image classification.

## Convolution Layer

Convolutional layers extract features from the input image. A convolutional layer’s units are arranged in feature maps. Each unit is related to local patches in the preceding layer’s feature maps through a collection of weights known as a filter bank. A feature map’s filter bank is shared by all units. Distinct filter banks are used by different feature maps in a layer. There are two reasons behind this design. First, with array data such as photos, local clusters of values are frequently highly linked, resulting in clearly detectable local themes. Second, image and other signal local statistics are unaffected by their position. In other words, if a motif can exist in one section of the picture, it may appear everywhere, which is why units in various places are given the same weights, and the same pattern is detected in other portions of the array [69].

### Activation Function

The primary function of activation is to remove redundant data and retain features extracted by convolutional layers, and map features using non-linear functions [70]. Many activation functions are used, such as ReLU [71], sigmoid [72], tanh [73] and Softmax [74].

### Pooling Layer

After the convolution layer, the pooling layer is usually applied. The primary role or task of the pooling layer is to shrink feature maps and compress the received feature map. The pooling processes include max pooling and average pooling [75, 76].

### Fully Connected Layers

Every CNN model eventually contains fully connected layers. Each cell in the layer connects to all the previous neurons. The fully connected layers are used as a classifier for CNN Model [77].

## Proposed METHODOLOGY

In this section, the PRCnet model, preprocessing and data set is discussed. The PRCnet model consists five blocks of convolutional layers. Each convolutional layer is followed by the ReLU activation function and batch normalization. The third, fourth and fifth blocks consist of three parallel convolutional layers with different filter sizes. There are six residual connections between layers. These connections for a better representation of features, obtaining features from different levels.

### Data sets

The first database (Dataset A) we used is the brain tumor dataset, introduced by Cheng et al. [51, 78]. It has 3064 MRI images of the brain collected from 233 patients in different views, including sagittal, coronal, and axial. It is classified into three classes according to the types of brain tumors . It contains 708 images of meningioma, 930 images of pituitary tumor and 1426 images of glioma. Images are available in Mat format and size 512 x 512 . The second dataset (Dataset B) is Brain Tumor MRI Dataset collected from the Kaggle website [79]. It contains 7023 labelled images grouped into four categories: 1621 images of glioma tumors, 1645 images of meningioma tumors, 2000 images of no tumor, and 1757 images of a pituitary tumor. Figure 4 shows the distribution of brain tumor categories for both datasets.Moreover, figure 5 presented a sample from the dataset that contains four categories of MRI.

**Fig 4.**
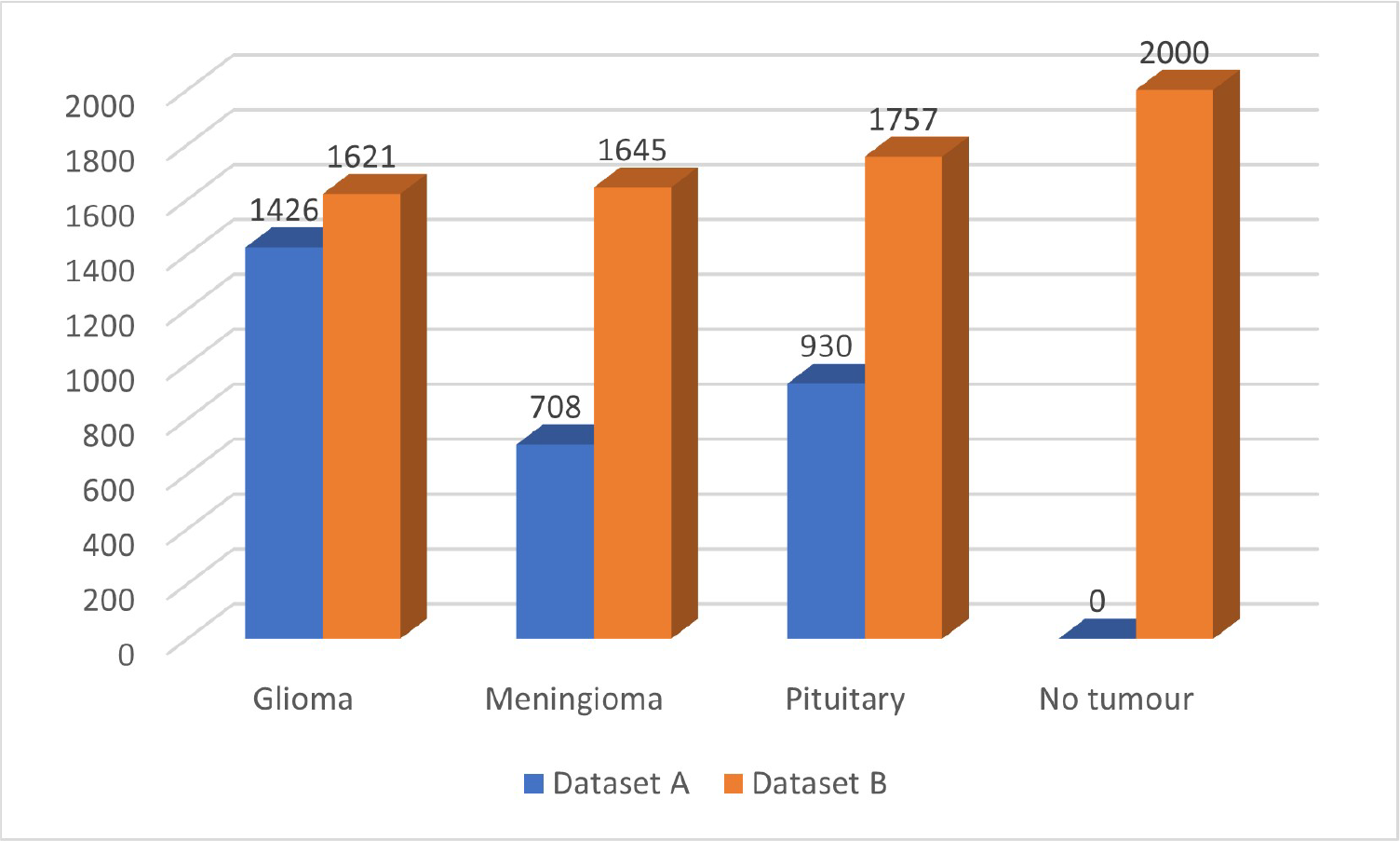
Distribution of brain tumor categories.

**Fig 5.**
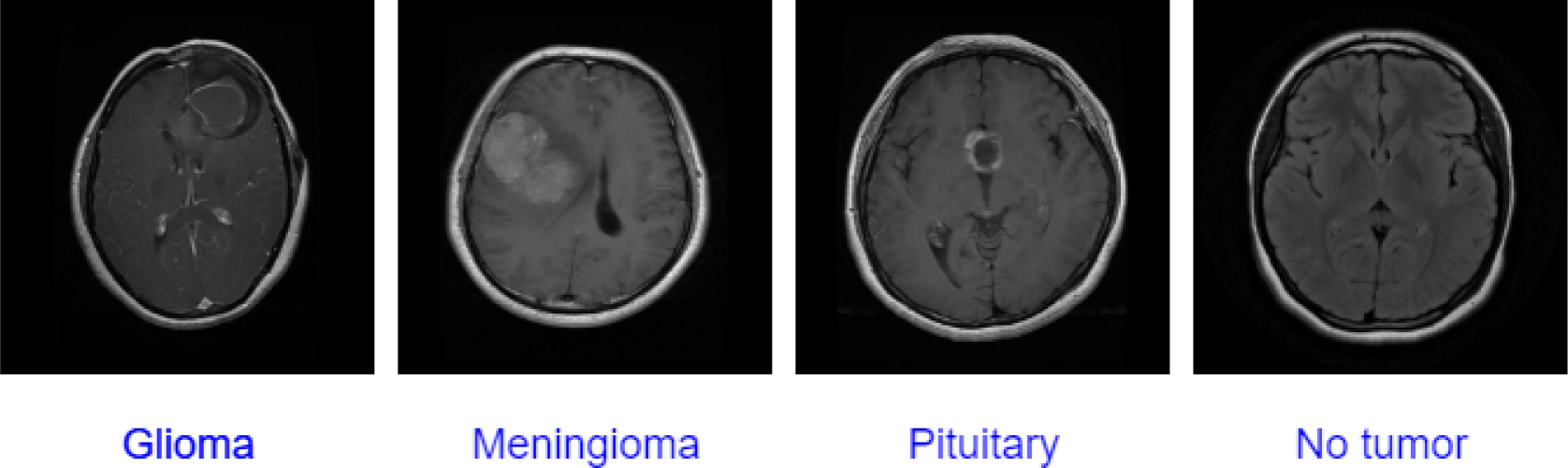
A Sample from the dataset B that presented the types of tumors.

### Pre-processing

All images in both data sets are resized to 224 x 224 to fit for the PRCnet, VGG16, VGG19 and mobileNetV2 input constraint. Then dataset A is divided into three groups 70% of the images are for training, 15% is validation, and the remaining 15% are for testing. On the other hand, dataset B is divided into three groups also 80% for training, 10% for the validation and 10% for the testing.

We used data augmentation techniques to increase the training dataset to overcome the lack of data for training. Many augmentation methods increase the training data set, which helps overcome the over-fitting problem. The augmentation techniques used in this paper are brightness, rotation, horizontal flip, vertical flip and shift.

### The PRCnet Model

This paper proposes an efficient CNN architecture to detect and classify brain tumors in MRI images.The PRCnet model used different filter sizes with parallel convolution layers also residual connections. The PRCnet model architecture mainly consists of convolution layers, max pooling, batch normalization and a fully connected layer. We chose a hyperparameter for the PRCnet model through several experiments. The optimal one was chosen after a random search was conducted using Keras Tuner on different sets of hyperparameters. Then, the best hyperparameters results chosen by the random search were compared with what was selected manually. Table 2 shows the values of the hyperparameters and the values that were chosen. Table 3 shows hyperparameter values and results obtained for the best eight models out of 20 trials, and each trial was tested twice. Also, table 3 shows the results obtained through manual hyperparameter values selection. The test was conducted on dataset B which is divided into two groups 80% for training and 20% for validation. The Batch size is set to 16, and the Epoch size is 30.

**Table 2.** Hyperparameters search space and which value is selected for PCRnet model.

| Hyperparameter | Search space | Value chosen |
| --- | --- | --- |
| No filters in Conv 1 | 32, 64 or 128 | 32 |
| Kernel size of Conv 1 | 3,5 or 7 | 3 |
| No filters in Conv 2 | 32, 64 or 128 | 64 |
| Kernel size of Conv 2 | 3,5 or 7 | 3 |
| No filters in Conv 3 | 32, 64 or 128 | 64 |
| Kernel size of Conv 3 | 3,5 or 7 | 3 |
| No filters in Conv 4 | 32, 64 or 128 | 64 |
| Kernel size of Conv 4 | 3,5 or 7 | 5 |
| No filters in Conv 5 | 32, 64 or 128 | 64 |
| Kernel size of Conv 5 | 3,5 or 7 | 7 |
| No filters in Conv 6 | 32, 64 or 128 | 128 |
| Kernel size of Conv 6 | 3,5 or 7 | 3 |
| No filters in Conv 7 | 32, 64 or 128 | 128 |
| Kernel size of Conv 7 | 3,5 or 7 | 3 |
| No filters in Conv 8 | 32, 64 or 128 | 128 |
| Kernel size of Conv 8 | 3,5 or 7 | 5 |
| No filters in Conv 9 | 32, 64 or 128 | 128 |
| Kernel size of Conv 9 | 3,5 or 7 | 7 |
| No filters in Conv 10 | 64, 128 or 256 | 256 |
| Kernel size of Conv 10 | 3,5 or 7 | 3 |
| No filters in Conv 11 | 64, 128 or 256 | 256 |
| Kernel size of Conv 11 | 3,5 or 7 | 3 |
| No filters in Conv 12 | 64, 128 or 256 | 256 |
| Kernel size of Conv 12 | 3,5 or 7 | 3 |
| No filters in Conv 13 | 64, 128 or 256 | 256 |
| Kernel size of Conv 13 | 3,5 or 7 | 3 |
| No filters in Conv 14 | 64, 128 or 256 | 256 |
| Kernel size of Conv 14 | 3,5 or 7 | 5 |
| No filters in Conv 15 | 64, 128 or 256 | 256 |
| Kernel size of Conv 15 | 3,5 or 7 | 7 |
| No. Units in Dense layer 1 | min value:256,max value:1024 | 256 |
| No. Units in Dense layer 2 | min value:256,max value:1024 | 256 |
| Learning rate | min value: 0.0001,max value:0.01 | 0.0001 |

**Table 3.** Hyperparameter values for the highest eight models accuracy results out of 20 trials as well as the results through manual hyperparameter values selection.

| Hyperparameter | Trial 1 | Trial 2 | Trial 3 | Trial 4 | Trial 5 | Trial 6 | Trial 7 | Trial 8 | Manual selection |
| --- | --- | --- | --- | --- | --- | --- | --- | --- | --- |
| No filters in Conv 1 | 32 | 128 | 128 | 32 | 128 | 32 | 64 | 32 | 32 |
| Kernel size of Conv 1 | 7 | 7 | 5 | 5 | 7 | 7 | 7 | 7 | 3 |
| No filters in Conv 2 | 64 | 64 | 32 | 32 | 128 | 64 | 64 | 64 | 64 |
| Kernel size of Conv 2 | 7 | 3 | 3 | 5 | 5 | 3 | 3 | 5 | 3 |
| No filters in Conv 3 | 64 | 128 | 128 | 32 | 64 | 128 | 128 | 64 | 64 |
| Kernel size of Conv 3 | 5 | 7 | 5 | 7 | 5 | 3 | 3 | 5 | 3 |
| No filters in Conv 4 | 128 | 64 | 32 | 32 | 32 | 128 | 64 | 32 | 64 |
| Kernel size of Conv 4 | 7 | 5 | 3 | 3 | 7 | 7 | 3 | 7 | 5 |
| No filters in Conv 5 | 128 | 128 | 64 | 128 | 128 | 32 | 64 | 64 | 64 |
| Kernel size of Conv 5 | 7 | 5 | 7 | 3 | 3 | 3 | 3 | 5 | 7 |
| No filters in Conv 6 | 128 | 256 | 64 | 256 | 256 | 64 | 128 | 256 | 128 |
| Kernel size of Conv 6 | 5 | 7 | 5 | 7 | 5 | 3 | 7 | 7 | 3 |
| No filters in Conv 7 | 64 | 128 | 256 | 128 | 64 | 64 | 256 | 64 | 128 |
| Kernel size of Conv 7 | 3 | 5 | 5 | 7 | 7 | 7 | 7 | 7 | 3 |
| No filters in Conv 8 | 256 | 64 | 256 | 64 | 64 | 256 | 256 | 64 | 128 |
| Kernel size of Conv 8 | 7 | 3 | 3 | 5 | 7 | 7 | 3 | 7 | 5 |
| No filters in Conv 9 | 128 | 256 | 128 | 128 | 256 | 128 | 256 | 256 | 128 |
| Kernel size of Conv 9 | 3 | 3 | 7 | 3 | 7 | 5 | 3 | 5 | 7 |
| No filters in Conv 10 | 256 | 64 | 128 | 64 | 64 | 64 | 256 | 64 | 256 |
| Kernel size of Conv 10 | 5 | 7 | 3 | 3 | 5 | 7 | 5 | 5 | 3 |
| No filters in Conv 11 | 256 | 256 | 128 | 128 | 128 | 128 | 256 | 128 | 256 |
| Kernel size of Conv 11 | 7 | 3 | 7 | 5 | 7 | 7 | 5 | 5 | 3 |
| No filters in Conv 12 | 128 | 64 | 64 | 64 | 128 | 64 | 128 | 64 | 256 |
| Kernel size of Conv 12 | 5 | 7 | 3 | 7 | 7 | 5 | 7 | 5 | 3 |
| No filters in Conv 13 | 64 | 64 | 64 | 256 | 256 | 64 | 256 | 64 | 256 |
| Kernel size of Conv 13 | 3 | 7 | 7 | 3 | 7 | 3 | 7 | 7 | 3 |
| No filters in Conv 14 | 256 | 256 | 256 | 64 | 128 | 256 | 128 | 64 | 256 |
| Kernel size of Conv 14 | 7 | 7 | 5 | 7 | 7 | 5 | 7 | 5 | 5 |
| No filters in Conv 15 | 64 | 256 | 256 | 64 | 64 | 64 | 256 | 256 | 256 |
| Kernel size of Conv 15 | 5 | 3 | 3 | 3 | 3 | 7 | 3 | 7 | 7 |
| No. Units in Dense layer 1 | 768 | 1024 | 256 | 1024 | 768 | 384 | 1024 | 640 | 256 |
| No. Units in Dense layer 2 | 640 | 384 | 256 | 384 | 896 | 256 | 640 | 512 | 256 |
| Learning rate | 0.001547 | 0.000535 | 0.002151 | 0.001764 | 0.001934 | 0.001197 | 0.000120 | 0.001566 | 0.0001 |
| Validation Accuracy | 0.8935 | 0.8787 | 0.8752 | 0.8718 | 0.8680 | 0.8618 | 0.8577 | 0.8390 | 0.9100 |

See figure 6 where the model is explained in detail. The size of the entered image is *224 x 224 x 1*. Starts a model with a convolutional layer with a filter size of *3 x 3)* and stride size of *2*. If we assume that the input image is I and the filter is K and its size *n x m*. The output of the convolution will be given by the equation [80].

**Fig 6.**
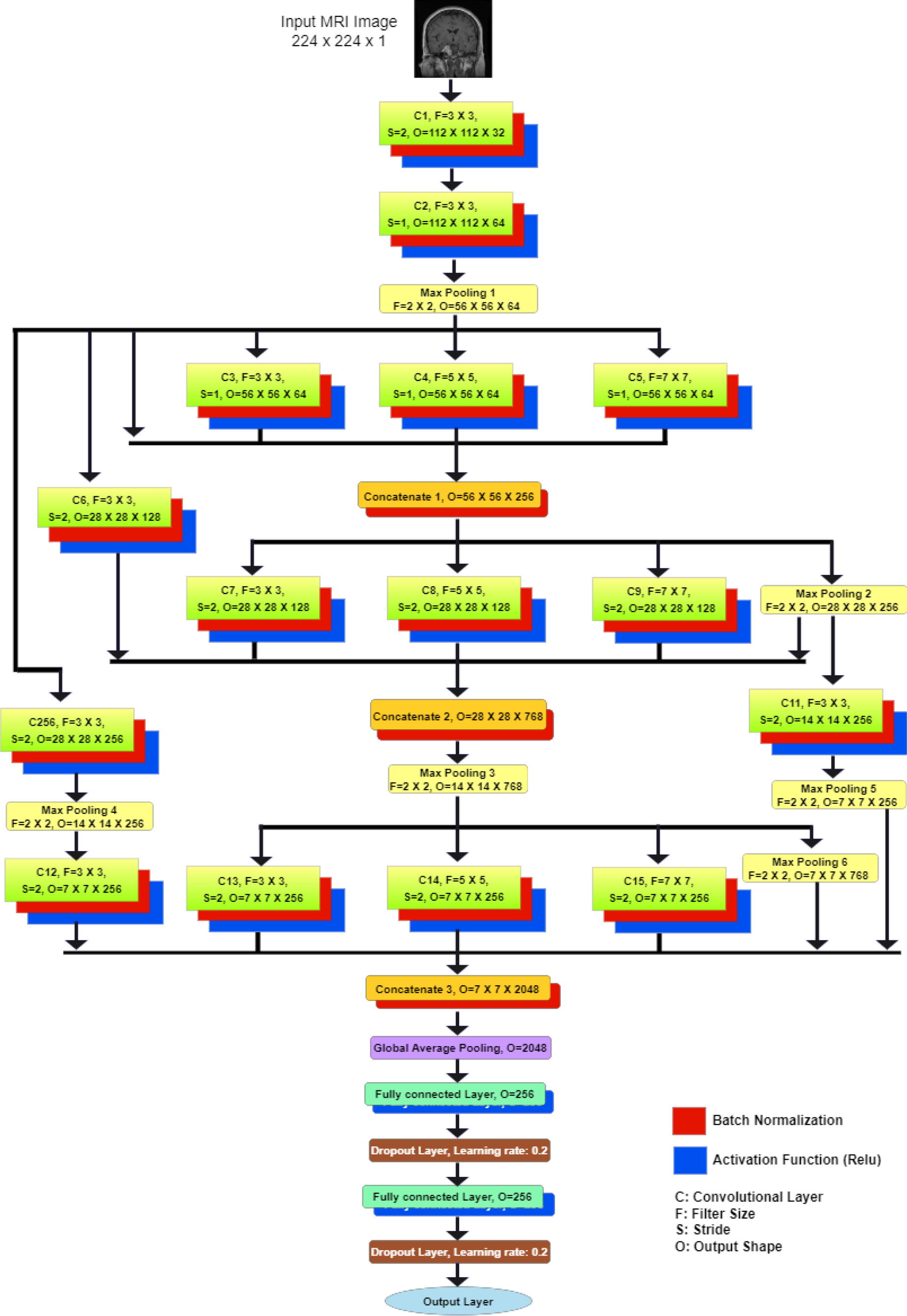
The PRCnet Model.

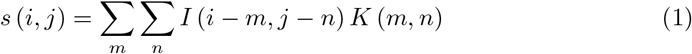

Where s is the output image, I is the input image, k is the kernel filter and n,m is the size of the kernal filter.

Each convolutional layer is followed by a batch normalization layer for speeding up the training process. Whereas, the batch normalization layer normalizes the input for each small batch and propagates the gradients through normalization parameters, as shown in equation [81].

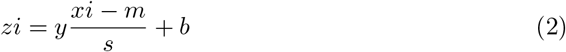

Here *xi* is the input value, *y* and b learned parameter, *m* is the batch mean, s is the batch standard deviation. The Relu activation function will be applied after batch normalization to reduce the effect of the vanishing gradient problem. The first convolutional layer is followed by a second convolutional layer with a filter size of *3 x 3* and a stride size of *2*. The second layer follows a Max Pooling layer with a filter size of *2 x 2*. Then comes three blocks of parallel convolutional layers. Each block consists of three parallel convolutional layers with filter sizes *(3 x 3, 5 x 5, 7 x 7)* the use of different filter sizes to ensure that the model recognizes small and large features. The stride size in the first block is one, and in the second and third block is *2*. These three layers are combined with the connections from the previous layers with the concatenation layer followed by batch normalization and the Max Pooling layer after the second block. There are six connections between layers, some of which are short and direct, and some of the connections go through convolutional layers or Max Pooling or both. These connections are for a better representation of features, obtaining features from different levels, and addressing the problem of gradient vanishing. After that, global average pooling will be applied to reduce the input dimensions size to one dimension. Finally, there are three fully connected layers, the first and second layers with the ReLU activation function, and the last layer is for the classification with a Softmax activation function. After the first and second fully connected layers, a dropout layer with a ratio of 0.2. Dropout layer help to avoid the over fitting problem.

## Experiment Results

In this section, we will present the experimental setup, adopted hyperparameters, evaluation metrics, and the results of testing the PRCnet model on the two datasets and comparing our results with other models.

## Experimental Setup

To implement the proposed model, we used the python 3.8.0 programming language, Keras 2.6.0, and Tensorflow 2.6.0 libraries. Specifications of the computer that was used is: Intel(R) Core(TM) i7-10750H CPU @ 2.60GHz 2.59 GHz, 16GB RAM, NVIDIA GeForce GTX 1660 Ti GPU and Windows 10 installed.

## Hyperparameters

Hyperparameters are the basic things that must be determined before starting the training process, including Batch size, Epoch size, and the learning rate, where the Batch size is set to 64, Epoch size is 250, and the learning rate is 0.0001. Moreover, the early stop was set to stop the training if there is no improvement in the validation Loss after 40 Epoch and store the weights that achieved minimum validation Loss.

## Evaluation metrics

Performance measures play an essential role in testing and evaluation in developing any machine learning model [82]. Furthermore, evaluation metrics are essential for measuring the accuracy of any classifier, as the results of any classifier may be good against specific metrics and may not be good or bad against other metrics [83]. Therefore, metrics should be used in the training and testing phases to evaluate the system. Here are some common metrics used to evaluate the system [84, 85].

1. **Accuracy**: The ratio of correct predictions is divided by the number of predictions evaluated.

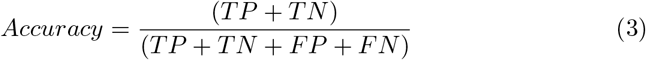
2. **Recall**: It is used to measure positive patterns that have been correctly categorized as in the following equation.

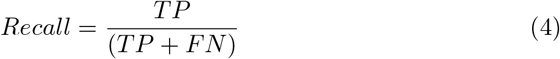
3. **Precision**: It is used to measure the proportion of positive patterns correctly predicted from the sum of the expected patterns in the positive category only.

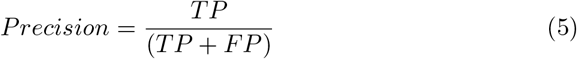
4. **F1-Score**: It represents the harmonic mean between the recall and accuracy values.

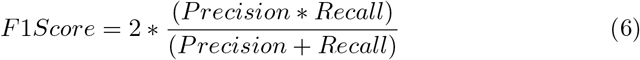

where,*TN* = True Negative, *FN* = False Negative, *TP* = True Positive, and *FP* = False Positive.

## Results

In this experiment, we trained and tested the PRCnet model on dataset A and dataset B and compared the results with the models (VGG16 [36], VGG19 [36], MobileNetV2 [68], Aurna et al [33], Chattopadhyay et al [37], Musallam et al [35]). As a result, the accuracy achieved by the PRCnet model was 94.77 on the dataset A and 97.1 on the dataset B, the details of dataset A results show in Table 4 and figure 9 while Table 5 and figure 10 show the results on dataset B. Also, we note that the result achieved by the stander models (VGG16, VGG19 and MobileNetV2) with Fine-Tuning is better than the accuracy without Fine-Tuning. These results are due to the cause these models are deep and need more data for training, so the results were better with fine tuning. The PRCnet model achieved a better result than the other models for several reasons. First, the standard models (VGG16, VGG19 and MobileNetV2) require a large amount of data for training. Therefore, we used transfer learning from ImageNet dataset, but the results not to the required level, due to the significant difference between the content of ImageNet dataset images and MRI images of the brain. Second, other CNN models such as (Chattopadhyay et al [37]) model is very simple and contains only two convolutional layers, which is not enough to extract all the features accurately, so the accuracy of this model was less compared to other models, especially as the size of the dataset increased. Also, the (Musallam et al [35]) model is complex and the computational cost is high because the kernel size for all convolutional layers is 7, so the accuracy of this model on dataset A was less than the accuracy of dataset B. Finally, the PRCnet model used parallel layers with filters of different sizes to ensure that the model learns the small and large features. In addition, the connection between different layers was used to help to better represent features from different levels and addresses the issue of gradient vanishing.

**Table 4.** Results on the Dataset A.

| Models |  | Accuracy (%) | Precision (%) | Recall (%) | F1-Score (%) |
| --- | --- | --- | --- | --- | --- |
| VGG16 | Without Fine-Tuning | 87.8 | 87.8 | 87.8 | 87.77 |
|  | With Fine-Tuning | 92.16 | 92.21 | 92.16 | 92.1 |
| VGG19 | Without Fine-Tuning | 88.24 | 88.2 | 88.24 | 88.2 |
|  | With Fine-Tuning | 88.24 | 89.19 | 88.24 | 88.52 |
| MobileNetV2 | Without Fine-Tuning | 90.2 | 90.01 | 90.2 | 90.08 |
|  | With Fine-Tuning | 90.85 | 90.99 | 90.85 | 90.91 |
| Aurna et al [33] |  | 92.59 | 92.54 | 92.59 | 92.47 |
| Chattopadhyay et al [37] |  | 83.44 | 83.18 | 83.44 | 83.17 |
| Musallam et al [35] |  | 80.39 | 79.66 | 80.39 | 79.94 |
| **The PRCnet Model** |  | **94.77** | **94.73** | **94.77** | **94.74** |

**Table 5.**
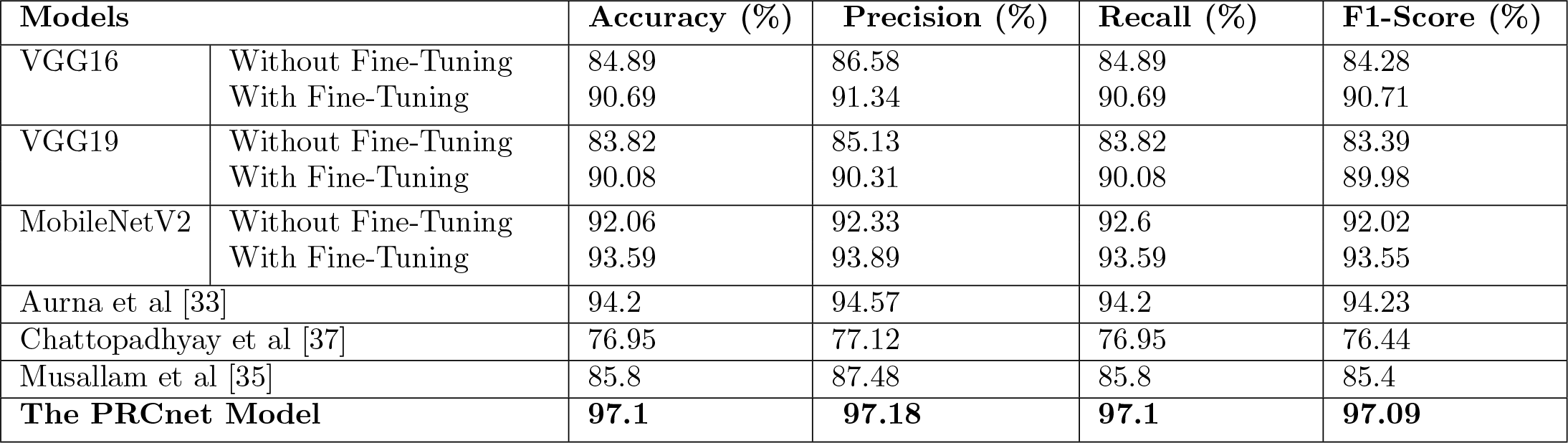
Results on the dataset B.

**Table 6.** Performance of the PRCnet model after applying stratified K-Fold cross-validation.

| Dataset | Fold | Accuracy (%) | Precision (%) | Recall (%) | F1-Score (%) |
| --- | --- | --- | --- | --- | --- |
| Dataset A | Fold 1 | 93.64 | 93.76 | 93.64 | 93.68 |
|  | Fold 2 | 95.76 | 96.09 | 95.76 | 95.82 |
|  | Fold 3 | 94.45 | 94.68 | 94.45 | 94.52 |
|  | Fold 4 | 94.94 | 94.92 | 94.94 | 94.93 |
|  | Fold 5 | 91.18 | 91.47 | 91.18 | 91.27 |
| Dataset B | Fold 1 | 95.73 | 95.80 | 95.73 | 95.67 |
|  | Fold 2 | 96.30 | 96.31 | 96.30 | 96.29 |
|  | Fold 3 | 96.65 | 96.71 | 96.65 | 96.66 |
|  | Fold 4 | 97.72 | 97.72 | 97.72 | 97.72 |
|  | Fold 5 | 96.65 | 96.71 | 96.65 | 96.60 |

Since the first and second datasets contain more than two classes, the Precision, Recall and F1-Score was made with average = weighted.

All models losses and accuracy of all models during the training and verification process on the dataset A are evident in figure 7 and on the dataset B shown in figure 8.

**Fig 7.**
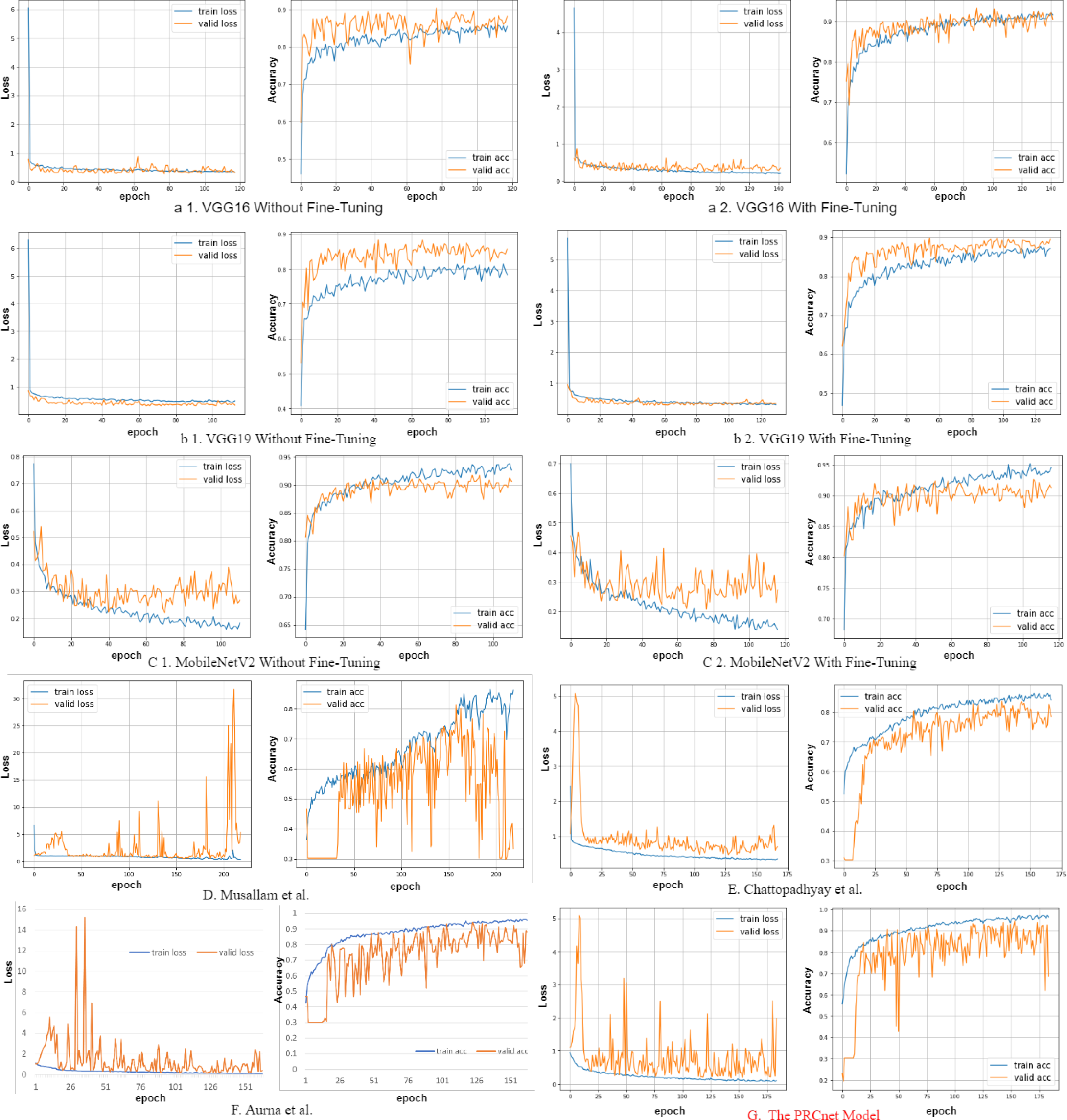
losses and accuracy of all models during the training and validation process on the dataset A.

**Fig 8.**
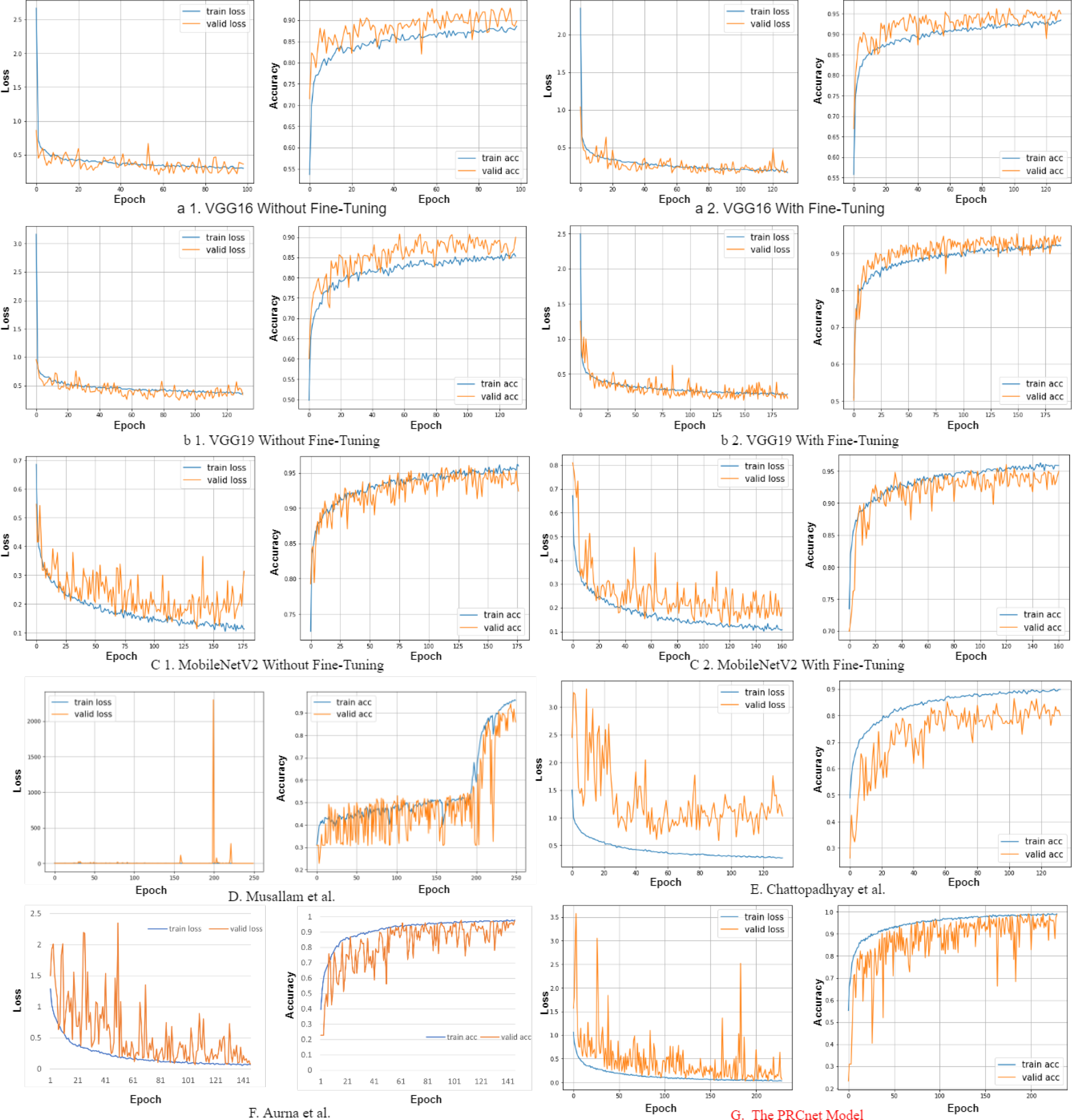
losses and accuracy of all models during the training and validation process on the dataset B.

**Fig 9.**
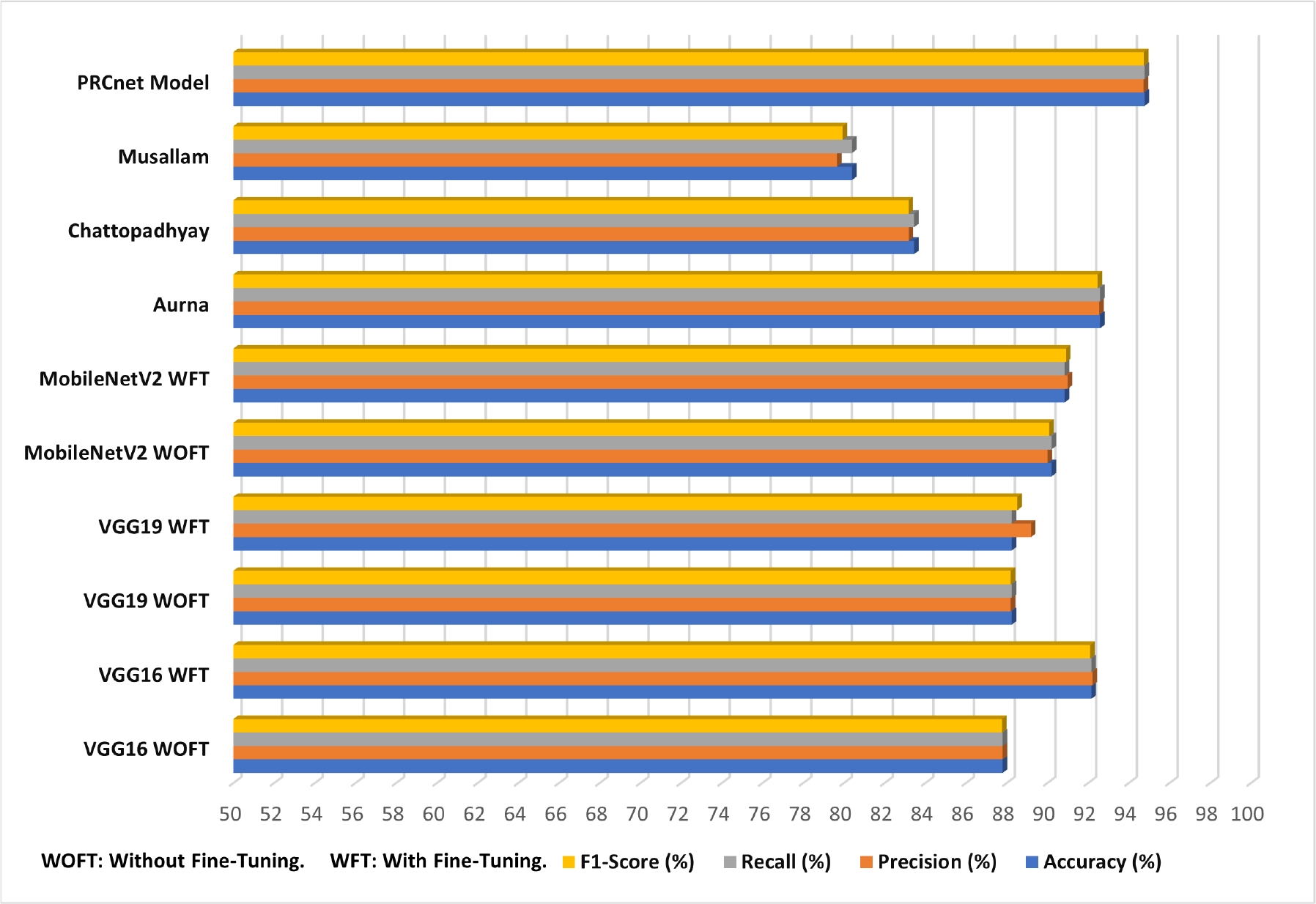
Results on the dataset A for all Models.

**Fig 10.**
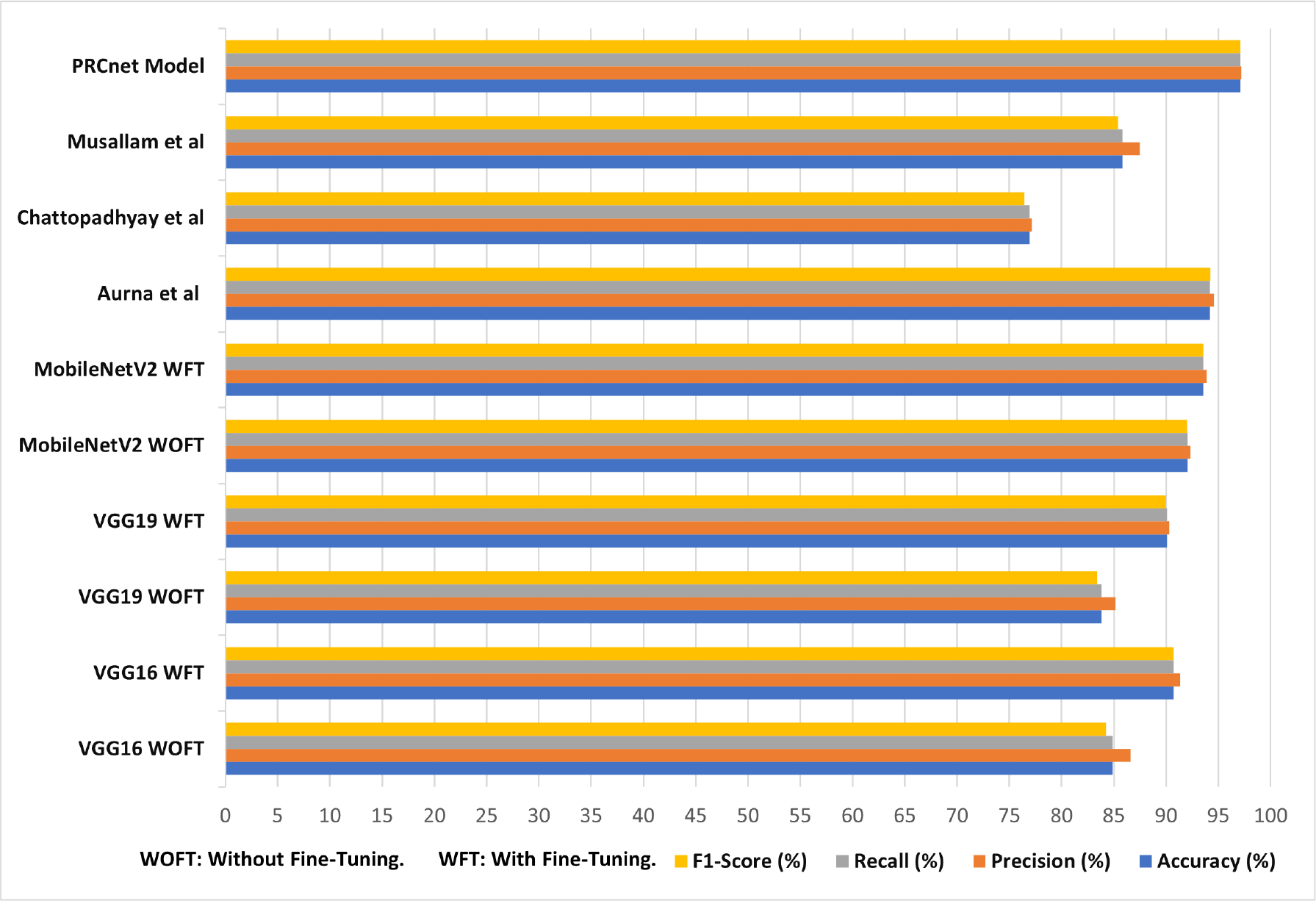
Results on the dataset B for all Models.

Figure 11 shows the confusion matrix for all models during the testing process on the dataset A. The results showed that the PRCnet model correctly identified 207 gliomas out of 214 images, 93 images of meningioma tumors out of 106 images, and detected 135 images of pituitary tumors correctly out of 139. The results during the testing process on the dataset B, as shown in figure 12, showed that the PRCnet model correctly distinguished all 150 pituitary tumors, correctly detected 140 out of 150 images of glioma, and correctly detected 146 images of meningioma out of 153 Picture and distinguish 200 images that are no-tumor out of 202. In general, it was noted that all models achieved the highest percentage of correct detection of pituitary tumors, no-tumors.

**Fig 11.**
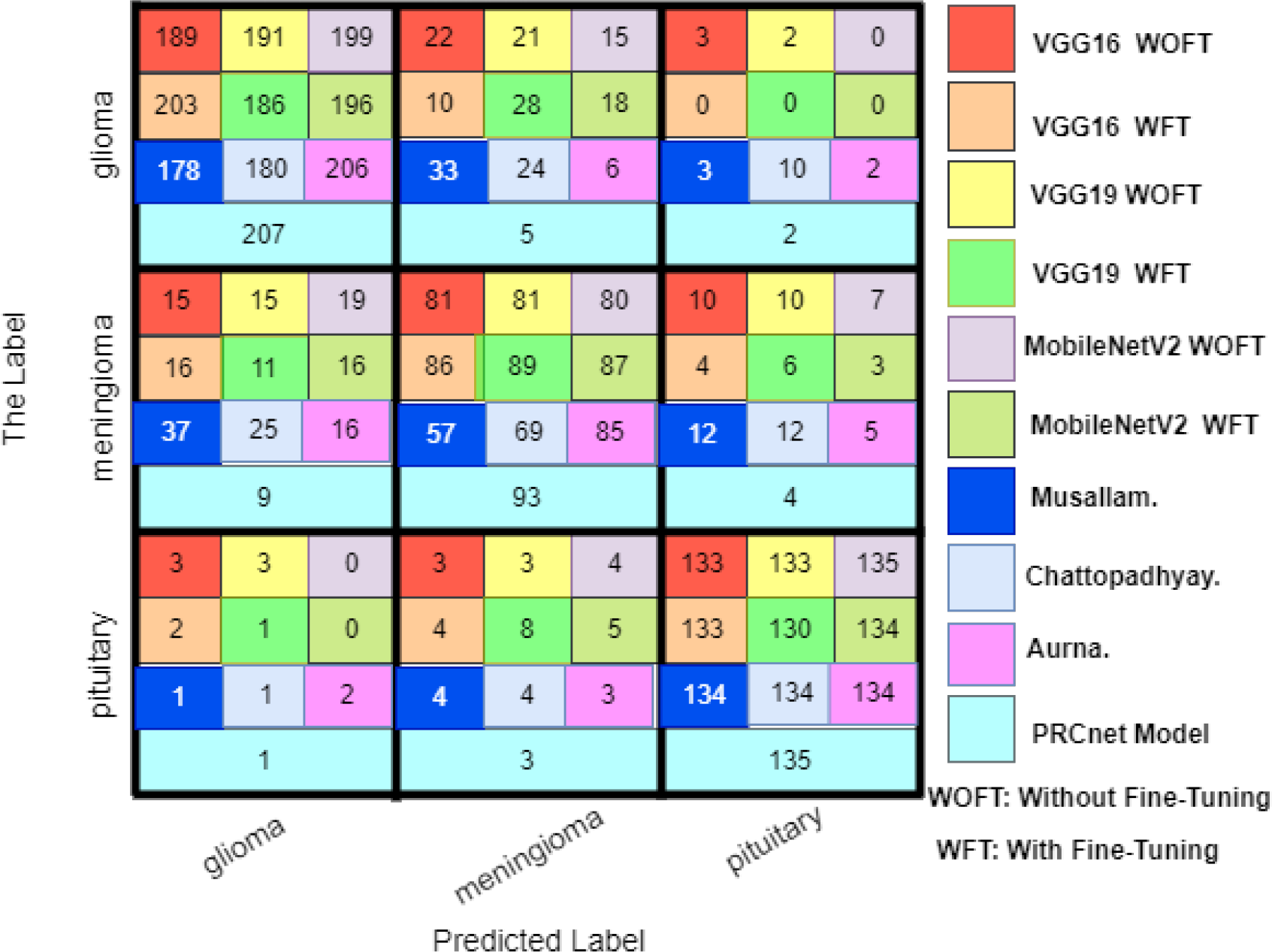
Confusion matrix for all models on the dataset A.

**Fig 12.**
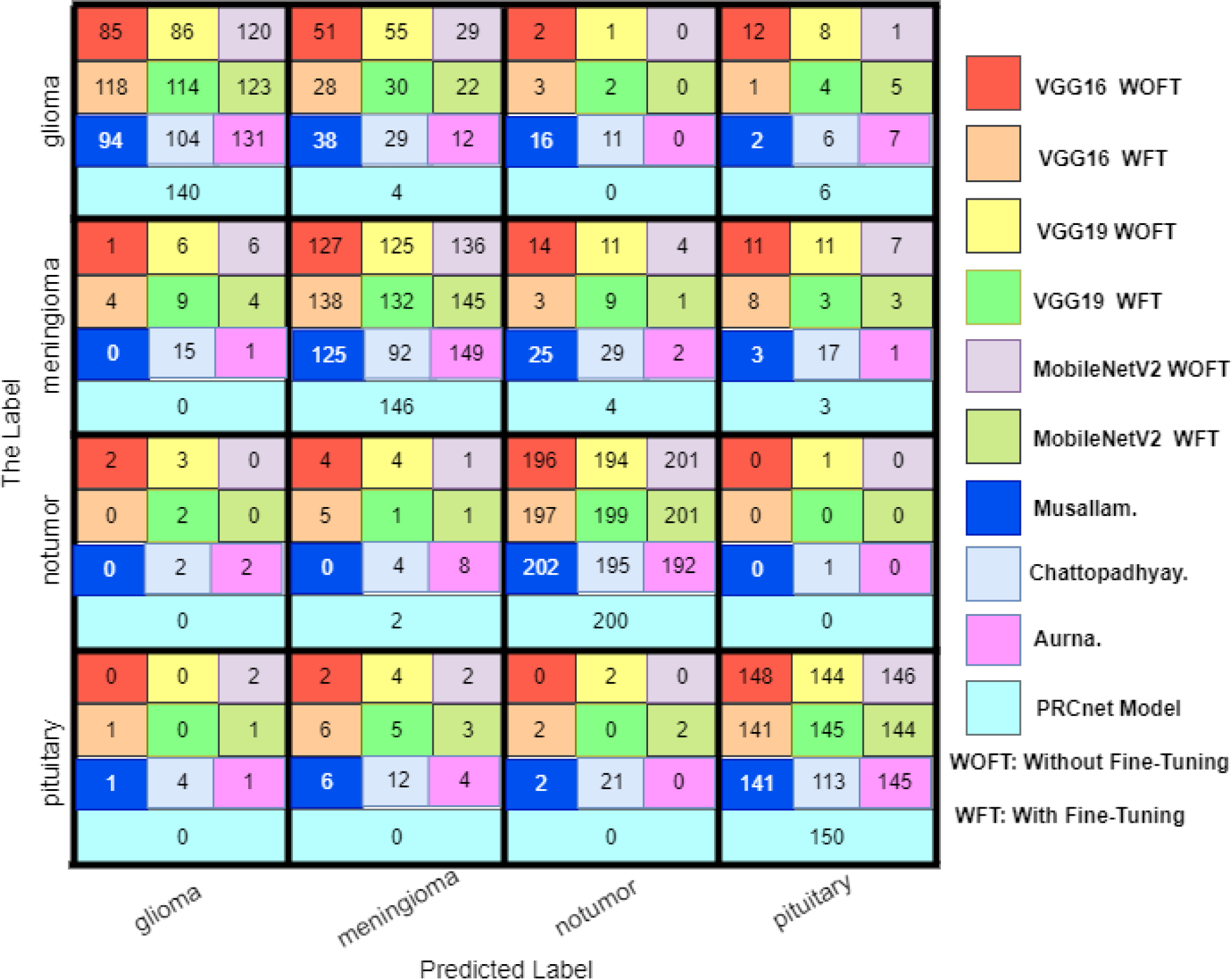
Confusion matrix for all models on the dataset B.

In addition to that, we used a Grad-CAM visualization technology [86] in the PRCnet model, as it was used on the last convolutional layer in the PRCnet model to create the heat map. As is known, the last convolutional layer shows the visualization description of the object in the model. Figure 13 illustrates the Grad-CAM visualization technology on the PRCnet models, where some random images were chosen. Moreover, Figure 14 shows the effect of convolutional layers on the image, where only six filters visualization are displayed for each convolutional layer.

**Fig 13.**
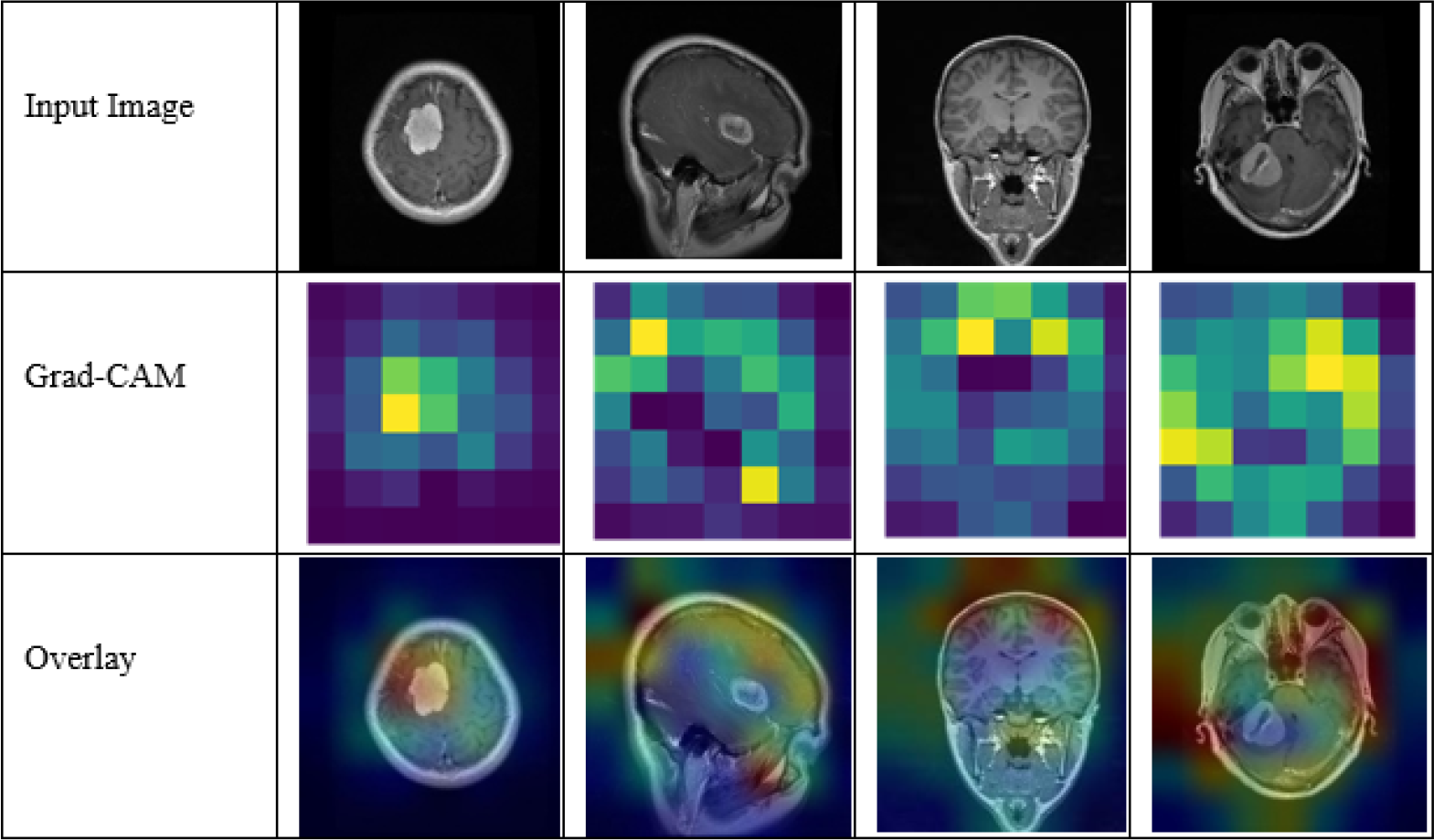
Grad-CAM visualization of features using PRCnet model.

**Fig 14.**
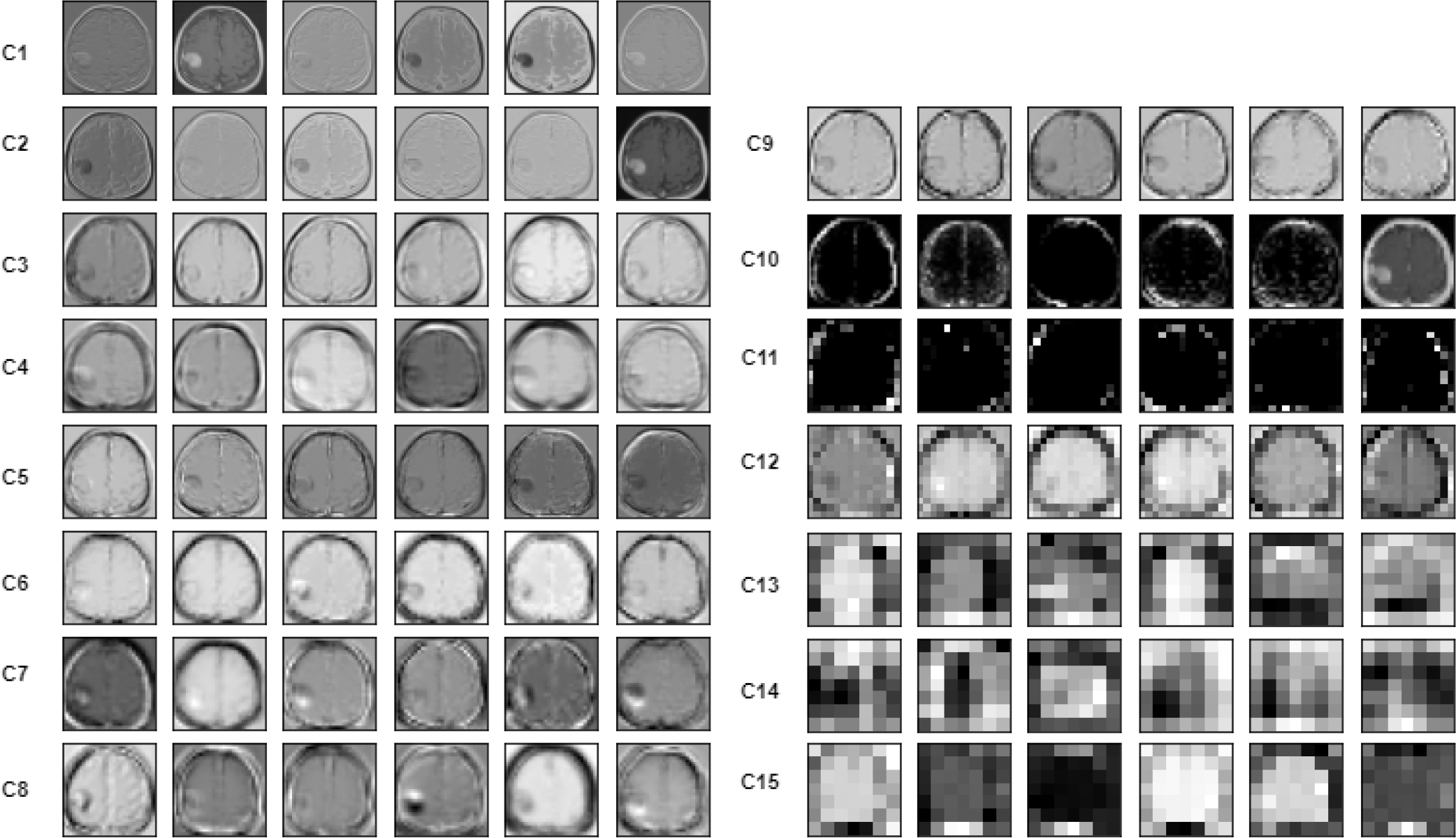
Samples visualization of six filters from each convolutional layer, C: Convolutional Layer.

### Stratified K-Fold cross validation

The cross-validation technique was used to verify the actual performance of the model and prove its robustness. The stratified K-Fold cross-validation method was used to validate the model. Where the data set is divided into k equal parts, the number of these parts depends on the size of the dataset [87, 88]. In this study, we determined the value of k = 5. After the data set has been divided into five parts, each part is considered a test set and the rest are considered training. Table **??** shows the results of testing the PRCnet model performance on datasets A and B using the stratified K-Fold cross-validation technique.

### The effect of data augmentation on the results

Developing a robust and reliable model for the detection of brain tumors requires a large and diverse data set in order to train it. Therefore, the application of deep learning in the field of medical images is rather difficult due to the lack of labeled training data [89]. In order to reduce the impact of lack of data in the proposed model, data augmentation was used, as five methods were used to increase the data as we mentioned in the pre-processing stage. The methods of augmentation were applied to the training data. The model was tested without the methods of augmentation and with the methods of augmentation, where the accuracy result was better with use the of data augmentation. Table 7 shows the accuracy of the model on both databases before and after using the data augmentation methods.

**Table 7.** The result before and after data augmentation.

| Dataset | Accuracy (%) Before data augmentation | Accuracy (%) After data augmentation |
| --- | --- | --- |
| Dataset B | 93.74 | 97.1 |
| Dataset A | 93.46 | 94.77 |

### The effect of convolution filter size on the results

In order to test the effect of filter size on the accuracy of the model, the model was tested on different filter sizes, as shown in the table 8. Experiments proved that the use of filters of different sizes gave better results because the use of different filter sizes helps the model to practice extracting large and small features.

**Table 8.** The result with a different filter size.

| Dataset | filter size | Accuracy (%) |
| --- | --- | --- |
| Dataset B | 3 x 3 | 92.37 |
|  | 5 x 5 | 96.49 |
|  | 7 x 7 | 96.49 |
|  | Multi filter size (3 x 3, 5 x 5, 7 x 7) | 97.1 |
| Dataset A | 3 x 3 | 91.5 |
|  | 5 x 5 | 91.72 |
|  | 7 x 7 | 93.46 |
|  | Multi filter size (3 x 3, 5 x 5, 7 x 7) | 94.77 |

## Discussion

In this study, we propose a PRCnet model for detecting brain tumors in MRI images. The PRCnet model uses parallel layers with different filter sizes to ensure that the model recognizes both small and large features in the MRI images. Additionally, the connection between different layers extracts features of different levels and helps overcome the gradient vanishing problem.

To evaluate the performance of the proposed PRCnet model, we conducted extensive experiments and comparative analyses on two datasets, namely Dataset A and Dataset B. The results of the experiments indicate that the proposed model achieved better results compared with the state-of-the-art models, with accuracy values of 94.77 and 97.1 for Dataset A and Dataset B, respectively.

We compared the results with the standard models, including VGG16, VGG19, and MobileNetV2, with and without fine-tuning. The accuracy of the standard models with fine-tuning on Dataset A is 92.16, 88.24, and 90.85 for VGG16, VGG19, and MobileNetV2, respectively. On the other hand, the accuracy of the standard models with fine-tuning on Dataset B is 90.69, 90.08, and 93.59 for VGG16, VGG19, and MobileNetV2, respectively. We used the transfer learning technique with standard models to obtain an initial weight from the ImageNet dataset and later fine-tuned it on the actual MRI image dataset. However, the proposed PRCnet model achieved better results than these standard models due to the deep of these models, which require a large dataset for training purposes. In addition, there is a significant difference between the ImageNet and MRI dataset content, negatively impacting the results.

We also compared the performance of the proposed model with the state-of-the-art models such as Aurna et al. [33], Chattopadhyay et al. [37], Musallam et al. [35].

Where accuracy achieved by the models Aurna, Chattopadhyay, and Musallam on Dataset A is 92.59, 83.44, and 80.39, respectively. While accuracy achieved on Dataset B is 94.2 for Aurna, 76.95 for Chattopadhyay, and 85.8 for Musallam. The Musallam and Chattopadhyay models achieved lower accuracy than Aurna and PCRnet models. The reason why The Chattopadhyay and Musallam model has lower accuracy compared to others is the model complexity. The Chattopadhyay model contains only two convolutional layers, which is insufficient to extract all features, leading to lower accuracy. On the other hand, the Muslim model is computationally costly due to the kernel size of 7 for all convolutional layers and requires more training datasets, which is reflected in the accuracy results for both Dataset A and Dataset B.

The proposed PRCnet model has achieved a reasonable classification rate, despite the limited dataset available for training and testing the model. However, there is still potential for further development of the model. For example, future research could explore new data augmentation methods to improve the model’s prediction ability.

Additionally, predicting the tumour’s location, grade, and size would be a valuable contribution to medical image analysis. This information could help physicians make more informed decisions about treatment options for their patients.

## Conclusion

This paper proposes the PRCnet model for classifying brain tumors using magnetic resonance imaging (MRI). We utilized several techniques, such as parallel convolutional networks with different filter sizes, to achieve accurate and automatic classification to improve feature representation. In addition, connections between different layers enabled us to obtain features at different levels, overcoming the problem of gradient vanishing. We also employed global average pooling and dropout between the fully connected layer to mitigate overfitting issues. The PRCnet model was tested on two datasets, the results shows PRCnet significantly outperform the state-of-the-art models, achieving an accuracy of 97.1 on dataset B and 94.77 on dataset A respectively. These results demonstrate the efficiency and generalizability of the proposed model and its potential to aid in the accurate and speedy diagnosis of brain tumors. To further improve the performance and accuracy of the model, future work can focus on pre-training the model on larger datasets such as Imagenet and transfer learning. Additionally, data augmentation methods can aid in better training the model and addressing any data imbalance issues.

## Notes

### Competing Interest Statement

The authors have declared no competing interest.

